# Dynamic-parameter movement models reveal drivers of migratory pace in a soaring bird

**DOI:** 10.1101/465427

**Authors:** Joseph M. Eisaguirre, Marie Auger-Méthé, Christopher P. Barger, Stephen B. Lewis, Travis L. Booms, Greg A. Breed

**Affiliations:** Department of Biology & Wildlife, University of Alaska Fairbanks, Fairbanks, AK, USA; Department of Mathematics & Statistics, University of Alaska Fairbanks, Fairbanks, AK, USA; Department of Statistics, University of British Columbia, Vancouver, BC, Canada; Institute for the Oceans & Fisheries, University of British Columbia, Vancouver, BC, Canada; Alaska Department of Fish & Game, Fairbanks, AK, USA; United States Fish & Wildlife Service, Juneau, AK, USA; Institute of Arctic Biology, University of Alaska Fairbanks, Fairbanks, AK, USA

**Keywords:** Bayesian, correlated random walk, golden eagle, movement ecology, soaring flight

## Abstract

Long distance migration can increase lifetime fitness, but can be costly, incurring increased energetic expenses and higher mortality risks. Stopover and other en route behaviors allow animals to rest and replenish energy stores and avoid or mitigate other hazards during migration. Some animals, such as soaring birds, can subsidize the energetic costs of migration by extracting energy from flowing air. However, it is unclear how these energy sources affect or interact with behavioral processes and stopover in long-distance soaring migrants. To understand these behaviors and the effects of processes that might enhance use of flight subsidies, we developed a flexible mechanistic model to predict how flight subsidies drive migrant behavior and movement processes. The novel modelling framework incorporated time-varying parameters informed by environmental covariates to characterize a continuous range of behaviors during migration. This model framework was fit to GPS satellite telemetry data collected from a large soaring and opportunist foraging bird, the golden eagle (*Aquila chrysaetos*), during migration in western North America. Fitted dynamic model parameters revealed a clear circadian rhythm in eagle movement and behavior, which was directly related to thermal uplift. Behavioral budgets were complex, however, with evidence for a joint migrating/foraging behavior, resembling a slower paced fly-and-forage migration, which could facilitate efficient refueling while still ensuring migration progress. In previous work, ecological and foraging conditions are normally considered to be the key aspects of stopover location quality, but taxa that can tap energy sources from moving fluids to drive migratory locomotion, such as the golden eagle, may pace migration based on both foraging opportunities and available flight subsidies.

## Introduction

Long-distance migration can relax competition and permit use of seasonally available resources, helping many animals maximize lifetime fitness (Newton, 2008; Avgar et al., 2014). Those benefits, however, come at substantial costs, including greater vulnerability to predators, uncertain conditions, mechanical wear, elevated energy expenditure, and time (Alerstam and Hedenström, 1998; Clark and Butler, 1999; Hedenström, 2008; Newton, 2008; Avgar et al., 2014). As many migrant species cannot store sufficient energy for nonstop, long-distance migration, stopover evolved as a behavior for strategically resting and refueling en route (Gill, 2007).

Migrant species are adapted for utilizing either soaring or flapping flight, and the different flight modes can relate to stopover strategy (Hedenström, 1993; Gill, 2007). Generally, soaring flight is favorable for larger birds and flapping flight for smaller birds, though the partitioning of time for each flight mode during migration is dependent on the relative importance of time and energy (Hedenström, 1993; Duerr et al., 2015; Katzner et al., 2015; Miller et al., 2016). In theory, a bird minimizing migration duration (time minimizer) would be expected to fly with greater directional persistence and stronger directional bias relative to a bird minimizing energy expenditure (or maximizing net energy gain). Such net energy maximizers would be expected to take advantage of en route foraging opportunities and may divert or delay to replenish energy reserves. (Note that “energy minimization” has been used to describe this strategy (e.g., Alerstam, 2011; Miller et al., 2016), but we use “net energy maximization” for clarity.) If time is less important, a net energy maximizer is less restricted and can spend additional time seeking an energetically superior path; the emergent path would then be more tortuous with less directional bias toward the final destination at any given point along the route. Time minimization and net energy maximization strategies are not mutually exclusive, however, and the emergent strategy and behaviors in any given migrating individual lies along a continuum (Alerstam, 2011; Miller et al., 2016).

Obligate soaring migrants must also consider routes based on their energy landscape (Shamoun-Baranes et al., 2010), the energetic constraints of movement over space (Shepard et al., 2013), which also contributes to a migrant’s location along the behavioral continuum. While soaring migrants can stopover, their energy landscape is more complex; at least as important as foraging resources are meteorological conditions, which can be extremely dynamic and subsidize the energetic cost of flight directly via uplift (Pennycuick, 1971; Alerstam, 1979; Spaar and Bruderer, 1997; Gill, 2007; Duerr et al., 2012; Murgatroyd et al., 2018). Two primary forms of uplift arise by (1) wind interacting with topography to form upslope wind or mountain waves (air currents forming standing waves established on the lee side of mountains; hereafter orographic uplift) and (2) solar heating of the earth’s surface to generate thermal uplift. Other forms arise from turbulent eddies over small landscape features and ocean waves modifying the air. The flight performance of soaring migrants relative to meteorological conditions has been well documented (Pennycuick, 1971; Alerstam, 1979; Spaar and Bruderer, 1997), establishing a clear link between diurnal migrant behavior and development of the atmospheric boundary layer. The dynamic nature of the atmospherically-driven flight subsidies requires detailed movement data as well as carefully designed analytical techniques to investigate certain mechanisms hidden in that data.

Our understanding of migratory processes has advanced enormously in the past 30 years, as animal tracking technology developed from a novelty of coarse observation to a core method for observing animal behavior and movement in incredible detail (Luschi et al., 1998; Sawyer et al., 2005; Bridge et al., 2011; Katzner et al., 2015; Hooten et al., 2017). Global Positioning System (GPS) telemetry, in particular, allows remote observation of animal relocations across a broad spatiotemporal scope. GPS transmitters are now light and reliable enough to study the complete migrations of many large soaring migrants, including golden eagles *Aquila chrysaetos*, which often rely on flight subsidies during migration (Katzner et al., 2015). Golden eagles and other large soaring birds have been used as model systems for phenomenologically evaluating questions about migratory flight performance and migration strategies (*sensu* Duerr et al., 2012; Lanzone et al., 2012; Katzner et al., 2015; Vansteelant et al., 2015; Miller et al., 2016; Shamoun-Baranes et al., 2016; Rus et al., 2017). For example, Lanzone et al. (2012) and Katzner et al. (2015) found that golden eagles use both thermal and orographic uplift to subsidize migratory flight, although thermal soaring was often more efficient in long distance, directed flight (Duerr et al., 2012). While these studies have contributed to our understanding of soaring migration and have laid a foundation for more detailed approaches, they relate meteorology to derived movement metrics, rather than with a process-based model to the movement data themselves, and ignore the temporal dependence between serially observed locations (i.e. autocorrelation), which imparts bias on certain estimated parameters (e.g., variances) thereby affecting inference through, for example, underestimating uncertainty. Consequently, the links between resources distributed over the landscape, such as flight subsidies, and behavioral budgets, including stopover behavior, during migrations of soaring birds remain unclear.

Unlike previous approaches, process-based, mechanistic movement models allow explicit inference of the underlying mechanisms driving movement (e.g., changes in behavior) that may not be available from conventional phenomenological analytical approaches (Turchin, 1998; Nathan et al., 2008; Hooten et al., 2017). While it is impossible to fully understand the intricacies in animal movement, we can pose mathematical models (e.g., correlated random walks) to approximate the movement process (Kareiva and Shigesada, 1983; Turchin, 1998), and then fit these models statistically to observed data to estimate parameters describing behavior and its relationship with dynamic environmental features moving animals experience (Blackwell, 1997, 2003; Morales et al., 2004; Breed et al., 2017; Hooten et al., 2017). Many of the recently developed mechanistic movement models are built in a discrete state-switching framework, where animals switch between discrete behavioral states (see Hooten et al., 2017, and references therein). Choosing both the biologically relevant and quantitatively supported number of states, as well as interpreting the identified states biologically remain challenging (Patterson et al., 2017; Pohle et al., 2017). Often, this challenge leads researchers to artificially limit the number of states and/or collapse two or more states into one biologically interpretable state. For example Pirotta et al. (2018), presented a model with five discrete kinds of avian flight, but the complexity of the model made interpreting those states biologically difficult and poorly matched classifications manually identified by an expert.

In many cases of animal movement, a more natural approach to modeling behavior is along a dynamic continuum, rather than as discrete states (Breed et al., 2012; Auger-Méthé et al., 2017; Jonsen et al., 2019). This may be especially the case for soaring birds, considering the inherently dynamic nature of atmospheric processes that influence movements. Here, we developed and applied a flexible mechanistic movement model based on a correlated random walk with time-varying parameters. This novel model was fit to movement data collected via GPS telemetry to understand how individuals in a population of long-distance soaring migrants use flight subsidies and budget stopover and migration behavior. Specifically, we were interested in identifying which flight subsidies influence stopover and migratory behavior and how the effect of key subsides and behaviors varied between spring and fall migrations. Our approach resembled continuous-time correlated random walks (Johnson et al., 2008; Blackwell et al., 2015; Gurarie et al., 2017; Michelot and Blackwell, 2019), but was easily implemented and estimated a relatively small number of dynamic parameters that could be directly interpreted biologically. A set of candidate models could be ranked using model selection approaches to infer how behavioral budgets and meteorological variables interacted to give rise to the observed migration paths. Modeling the effects of dynamic wind and uplift variables as time-varying further allowed new details in the behavior and movement of migratory golden eagles to emerge without imposing behavior to subjective discrete categorization.

## Methods

### Model system

Golden eagles are a large soaring raptor distributed across the Holarctic (Watson, 2010). While many populations are classified as partial migrant, most individuals that summer and breed above approximately 55°N in North America are considered true long-distance migrants (Watson, 2010; Kochert et al., 2002). Golden eagles are predatory and opportunistic, utilizing many taxa for food resources, ranging from small mammals and birds to ungulates, often scavenging carrion (Kochert et al., 2002; Watson, 2010). The population we observed in this study migrates over the mountainous regions of western North America between a breeding range primarily in southcentral Alaska, USA and a broad overwintering range in western North America that ranges from the southwestern US to central British Columbia and Alberta, Canada (Bedrosian et al., 2018).

### Data collection

We captured golden eagles with a remote-fired net launcher placed over carrion bait near Gunsight Mountain, Alaska (61.67°N 147.35°W). Captures occurred during spring, in mid-March to mid-April 2014-2016. A total of 53 adult and sub-adult eagles were equipped with 45-*g* back pack solar-powered Argos/GPS platform transmitter terminals (PTTs; Microwave Telemetry, Inc., Columbia, MD, USA). Eagles were sexed molecularly and aged by plumage.

PTTs were programmed to record GPS locations on duty cycles, ranging from 8-14 fixes per day during migration, depending on year of deployment. PTTs deployed in 2014 were set to record 13 locations at one-hour intervals centered around solar noon plus a location at midnight local time. PTTs deployed in 2015 were programmed to record 8 locations with one-hour intervals centered around solar noon, and PTTs deployed in 2016 took eight fixes daily at regular 3-hr time intervals. Note that the PTTs deployed in 2015 did not record locations overnight. Poor battery voltage from September to March often resulted in PTTs failing to take all programmed fixes, so the resulting GPS tracks had missing observations during these periods. Tags lasted multiple seasons and in fact many are still deployed and transmitting at this writing. For the analyses presented here, we analyzed the spring and fall migratory pathways of the 2016 migration from a sample of the tags deployed in 2014 (11), 2015 (7), and 2016 (8).

Movement data were managed in the online repository Movebank (https://www.movebank.org/), and we used the Track Annotation Service (Dodge et al., 2013) to extract flight subsidy (spatiotemporally explicit wind and uplift) data for each PTT location along eagle tracks. The Track Annotation Service derives uplift variables from elevation models and weather and atmospheric reanalyses (Bohrer et al., 2012). We followed the Movebank recommendations for interpolation methods; details are below.

### Movement model

We developed a correlated random walk (CRW) movement model to reveal how changes in behavior give rise to the movement paths of migrating eagles. Behavioral models that use a discrete-state switching framework have proven powerful and useful for answering a variety of ecological questions (e.g., Morales et al., 2004; Jonsen et al., 2005; Breed et al., 2009; Langrock et al., 2012). For this analysis, however, we chose to use a dynamic, time-varying correlation parameter, which represents behavior as a continuum rather than discrete categories, to capture complex behavioral patterns that could occur on multiple temporal and spatial scales (Breed et al., 2012; Auger-Méthé et al., 2017; Jonsen et al., 2019). We believe this approach can offer substantial flexibility, as a continuous range of behaviors is more realistic and, as we show, more naturally allows modeling behavior as a function of covariates.

The basic form of the model was a first-difference CRW presented by Auger-Méthé et al. (2017), which can take the form:

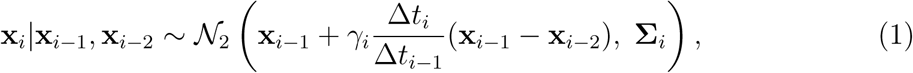

where

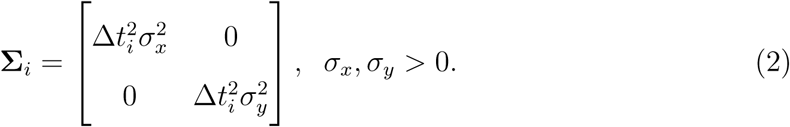

Here, Δ*t*_*i*_ = *t*_*i*_ − *t*_*i*−1_ represents the time interval between Cartesian coordinate vectors **x**_*i*_ and **x**_*i*−1_ for the observed locations of the animal at times *t*_*i*_ and *t*_*i*−1_. Incorporating autocorrelation in behavior, *γ*_*i*_ was a random walk, such that

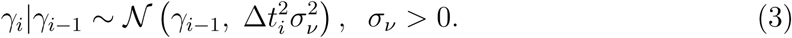

*γ*_*i*_ correlates displacements (or ‘steps’) and can be interpreted to understand the type of movement, and thus behavior, of migrating individuals: estimates of *γ*_*i*_ closer to one indicate directionally-persistent, larger-scale migratory movement, while estimates of *γ*_*i*_ closer to zero indicate more-tortuous, smaller-scale stopover movement (Breed et al., 2012; Auger-Méthé et al., 2017). Scaling *γ*_*i*_ by 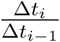 and the variance components by 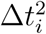 allows the model to accommodate unequal time intervals (Auger-Méthé et al., 2017), which can arise from a PTT’s pre-programmed duty cycles and/or missed location attempts. This assumes that over longer time intervals an animal is likely to move greater distances and that the previous step will have less influence on the current step. Notably, in introducing Δ*t*_*i*_, this CRW essentially becomes a correlated velocity model presented in terms of displacement vectors (**x**_*i*−1_ − **x**_*i*−2_) (Johnson et al., 2008; Blackwell et al., 2015; Gurarie et al., 2017), most closely resembling the autocorrelated velocity model presented by Gurarie et al. (2017). This model does not consider spatial error in observations. Given the scale of movements and precision of GPS technology, we decided it unnecessary to incorporate an observation equation, which would have increased model complexity. Fixing covariance to zero (equation 2) assumes that movement in the *x* and *y* dimensions are independent, and although we made this assumption, covariance could easily be accounted for at the cost of increased model complexity.

Extending this CRW to introduce environmental covariates, we first made the assumption that an individual’s behavior can be adequately explained by the previous behavior plus some effects of environmental conditions and random noise. This modeling approach and philosophy aligns with the movement ecology paradigm presented by Nathan et al. (2008): An animal’s movement path is influenced by its internal state and the environmental conditions it experiences. We modified the behavioral (or internal state) process—previously described above as a pure random walk in one dimension (equation 3)—similar to a linear model with a *logit* link function. The *logit* link constrains *γ*_*i*_ ∈ [0, 1] and allowed us to model it as a linear combination of continuously-distributed random variables (Jonsen et al., 2019). These variables were different meteorological conditions affecting flight subsidies. Now,

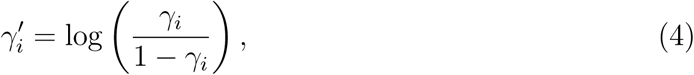

where

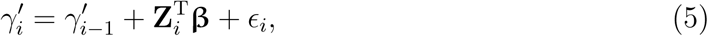

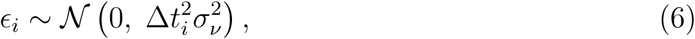

and 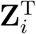 is the row vector of environmental covariates associated with **x**_*i*_. Each element of the vector **β** is an estimated parameter representing the magnitude and direction of the effect of its respective covariate on the correlation parameter *γ*_*i*_ in addition to the effect of *γ*_*i*−1_. Note that including *γ*_*i*−1_ here preserves explicit serial correlation in the behavioral process so that any additional environmental effect is not overestimated. 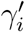 is only used to estimate *γ*_*i*_; any behavioral interpretations are made in terms of *γ*_*i*_.

### Model fitting

#### Subsetting tracks

To illustrate the application of how the presented modeling framework can provide new insight into animal movement ecology, we fit the model to 15 spring and 16 fall adult (i.e. entering their fifth or later summer) golden eagle migration tracks recorded by 18 adult males and 8 adult females in 2016. Note that this included repeated observations of migrations for five individuals; however, these observations were from different seasons (i.e., all observed fall migrations were from different individuals, as were all recorded spring migrations). In reporting the results, we assumed any effect of the few repeated measures to be negligible, which seems reasonable given fitted parameters presented in Table S1. The model was fit only to the migratory periods, plus two fixes prior to departure to ensure valid parameter estimates at the onset of migration. Data were subset to migratory periods under the following rules: The first migration step was identified as the first directed movement away from what was judged to be an individual’s summer (or winter) range with no subsequent return to that range, and the final migration step was defined as the step terminating in the apparent winter (or summer) range. This assignment was usually straightforward; however, in some cases there were apparent pre-migration staging areas. These were not considered part of migration and excluded from the analysis here.

#### Environmental covariates

Golden eagles can switch between using thermal and orographic uplift as flight subsidies (Lanzone et al., 2012; Katzner et al., 2015), so we included these variables as covariates affecting the correlation parameter in the behavioral process of the CRW (equation 5). Thermal uplift **z**_*tu*_ and orographic uplift **z**_*ou*_ are measured in m/s with **z**_*tu*_, **z**_*ou*_ ∈ [0, ∞). Thermal uplift was bilinearly interpolated from European Centre for Medium-Range Weather Forecasts (ECMWF) reanalyses, and orographic uplift from the nearest neighbor (grid cell) by pairing National Center for Environmental Predictions (NCEP) North American Regional Reanalysis (NARR) data with the Advanced Space-borne Thermal Emission Reflection Radiometer (ASTER) Global Digital Elevation Model (GDEM; Brandes and Ombalski, 2004; Bohrer et al., 2012). We also introduced wind as a covariate in the behavioral process, as it can influence eagle flight as well as the flight and energy landscape of many birds during migration (Shamoun-Baranes et al., 2017). Wind data were bilinearly interpolated from the NCEP NARR *u* (easterly/zonal) and *v* (northerly/meridional) components of wind predicted 30 m above ground, from which we calculated the tailwind support **z**_*tw*_, such that **z**_*tw*_ ∈ (−∞, ∞) (Safi et al., 2013), where positive values correspond to tailwind and negative values headwind. The bearings used to calculate each *z*_*tw,i*_ were the compass bearings required to arrive at **x**_*i*+1_ from **x**_*i*_.

We included a time of day interaction in the model because of clear diurnal effects. This also helped reduce zero inflation, particularly for thermal uplift, which often decays to zero after sunset due to heat flux and atmospheric boundary layer dynamics. To introduce the interaction, we used a dummy variable **z**_0_, such that *z*_0,*i*_ = 0 when *t*_*i*_ fell after sunset but before sunrise and *z*_0,*i*_ = 1 when *t*_*i*_ fell after sunrise but before sunset. This assumed behavior was not dependent on the covariates at night, which is sensible given observed diurnal behavioral cycles. Sunrise and sunset times local to each GPS point were calculated in R with the ‘sunriset’ function in the package ‘maptools’ (R Core Team, 2016; Bivand and Lewin-Koh, 2016). The final overall formulation of the behavioral process for the full model was:

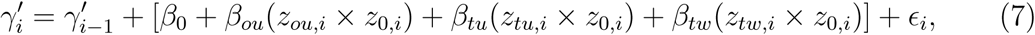

Prior to fitting the model, we followed Gelman et al. (2008) and *log*-transformed the uplift covariates and standardized variance to 0.25. We used a shifted *log*-transformation (Fox and Weisberg, 2019); adding one to the covariates prior to the *log*-transformation preserved zeros (i.e. zeros mapped to zero under the transformation). The distribution of raw tail wind support data appeared Gaussian, so it was only centered and standardized.

#### Parameter estimation & model selection

We fit our CRW in a Bayesian framework. Because the model has explicit serially correlated parameters, we used Hamiltonian Monte Carlo (HMC) over more conventional Markov-chain Monte Carlo (MCMC; e.g., Metropolis steps) to sample efficiently from a posterior with such correlation. Preliminary model testing indicated that HMC performed better, but note that a formal comparison of MCMC algorithms is beyond the scope of this paper.

Gelman et al. (2008) suggested Cauchy priors for logistic regression parameters; however, Ghosh et al. (2015) found that sampling from the posterior can be inefficient due to the fat tails of the Cauchy distribution. We thus chose Student-*t* priors with five degrees of freedom as weakly informative priors for the covariate parameters. Centering the prior density on zero allowed convergence to nonzero coefficient estimates only when such were sufficiently supported by the data. Weakly informative normal priors were placed on the variance parameters of the model.

We implemented HMC with R and Stan through the package ‘rstan’ (R Core Team, 2016; Stan Development Team, 2016). Working R and Stan code, including details on prior choice, and example data are provided as supplementary material, as well as supplementary tables and figures (Appendices 1-3). The model was fit to each track independently with five chains of 300,000 HMC iterations, including a 200,000 iteration warm-up phase, and retaining every tenth sample. Convergence to the posterior distribution was checked with trace plots, effective sample sizes, posterior plots of parameters, and Gelman diagnostics 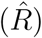 for each model fit.

We compared candidate models with leave-one-out cross-validation approximated by Pareto-smoothed importance sampling (PSIS-LOO) in R with the package ‘loo’ (Vehtari et al., 2016, 2017). The candidate models included possible combinations of environmental covariates plus a null CRW model without covariates. To limit model complexity and because we were interested in competing hypotheses about key predictors of behavior, we chose not to include interactions beyond time-of-day. We ranked the models by the expected *log* pointwise predictive density (i.e. out-of-sample predictive accuracy) transformed onto the deviance scale (looic; Vehtari et al., 2017), which allowed applying the rules of more traditional information-theoretic model selection (e.g., Burnham and Anderson, 2004). The model with the lowest looic was considered the best fit to the data, but if other models were within two looic of the top model, each, including the top model, were considered equally supported by the data.

## Results

### Model performance & diagnostics

Chain mixing, Gelman diagnostics 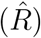 close to one, and large effective sample sizes for all parameters indicated convergence to the posterior of our CRW model for most model fits. Posteriors of parameters appeared Gaussian, also indicating convergence (Fig. S1). For five migration tracks, we did not consider the null model converged to the posterior (e.g., 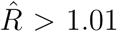), but in all cases the full model showed strong evidence of convergence. The five migrations for which the null model did not converge were excluded from formal model selection. Note that these five failed convergences were out of a total of 248 fits of the candidate model formulations across migration tracks.

### Behavior during migration

Median (interquartile range) departure and arrival dates were 5 March (4.5 *d*) and 27 March (6.4 *d*) in the spring and 29 September (11.7 *d*) and 16 November (15.5 *d*) in the fall. On average, eagles encountered similar orographic uplift in spring and fall but more intense thermal uplift and tailwind in the spring (Table 1).

**Table 1:**
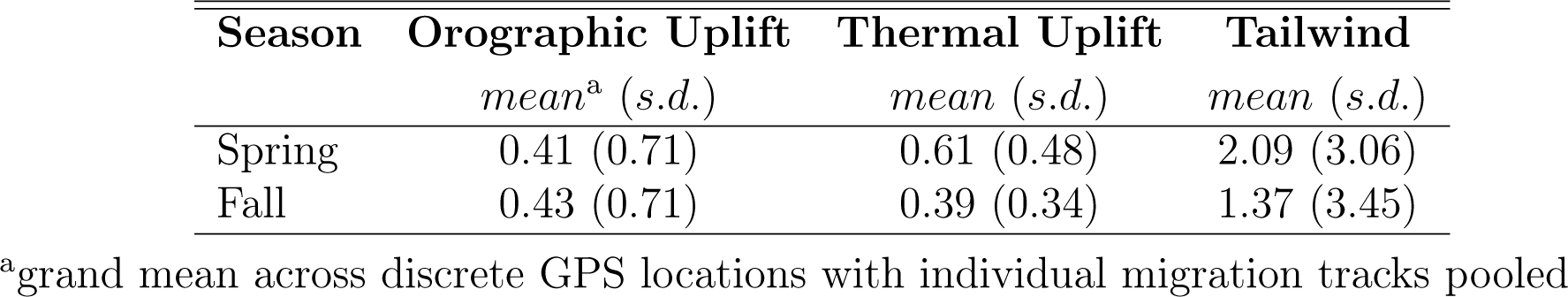
Summary statistics of flight subsidies encountered by migrating golden eagles that summer in Alaska. Variables were interpolated in space and time from weather reanalyses to eagle locations recorded by GPS telemetry. Units for all variables are *m/s*.

| **Season** | **Orographic Uplift** | **Thermal Uplift** | **Tailwind** |
| --- | --- | --- | --- |
|  | *mean<sup>a</sup> (s.d.)* | *mean (s.d.)* | *mean (s.d.)* |
| Spring | 0.41 (0.71) | 0.61 (0.48) | 2.09 (3.06) |
| Fall | 0.43 (0.71) | 0.39 (0.34) | 1.37 (3.45) |

<sup>a</sup>grand mean across discrete GPS locations with individual migration tracks pooled

The model revealed that eagles changed their behavior on multiple scales. First, there were very strong daily rhythms in behavior during migration, with birds migrating or moving more slowly and tortuously during the day and stopping at night (Figs. 1 & 2). While a time of day interaction explicitly in the model might drive a daily rhythm itself, accounting for serial correlation in behavior limited the overestimation of such an effect, as noted above and apparent in figure 2. Second, there was some evidence of stopover-like behavior, with individuals generally tending to continue moving along the migration route but exhibiting less directional persistence in movement (Fig. 3). The continuation along the migration route while in a stopover-like state is highlighted by track segments extended over space associated with low and intermediate estimates of *γ*_*i*_ (blue/purple in figure 3).

**Figure 1.**
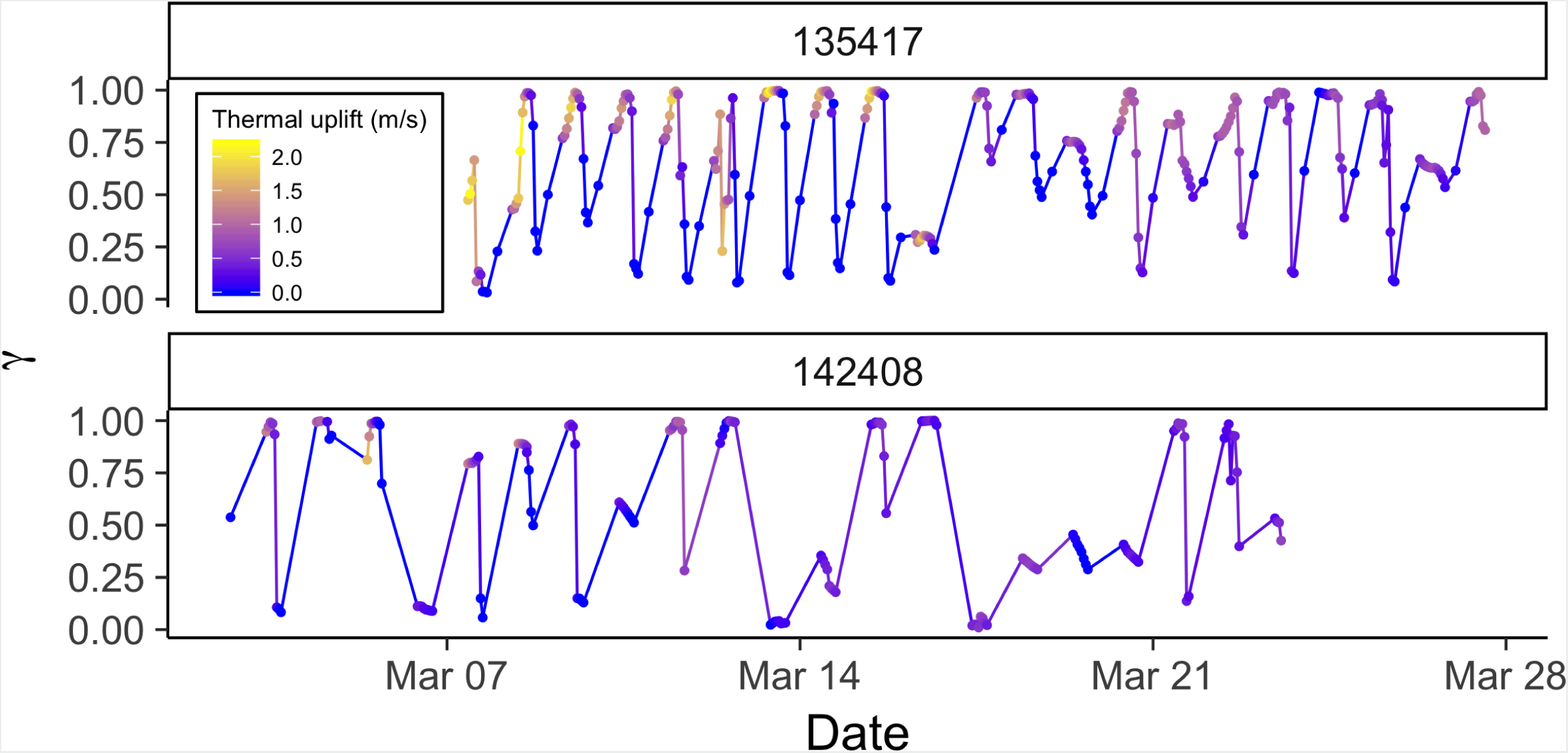
Time series of behavior parameter *γ* from correlated random walk model with full behavioral process (orographic uplift, thermal uplift, and tailwind support as predictors) for two golden eagles during spring migration with PTTs reporting on different duty cycles. Upper panel is 13 hourly centered on solar noon plus one at midnight, and the lower panel is 8 hourly centered on solar noon. *γ* close to one reflect movements associated with migratory behavior, and *γ* close to zero stopover behavior. Points are times of observations, and lines are linear interpolations between points. Hue indicates intensity of thermal uplift, with yellow indicating greater and blue lower. Note the daily rhythm in behavior associated with intense thermal uplift, stopover periods of one or more days, and the intermediate periods suggesting fly-and-forage.

**Figure 2.**
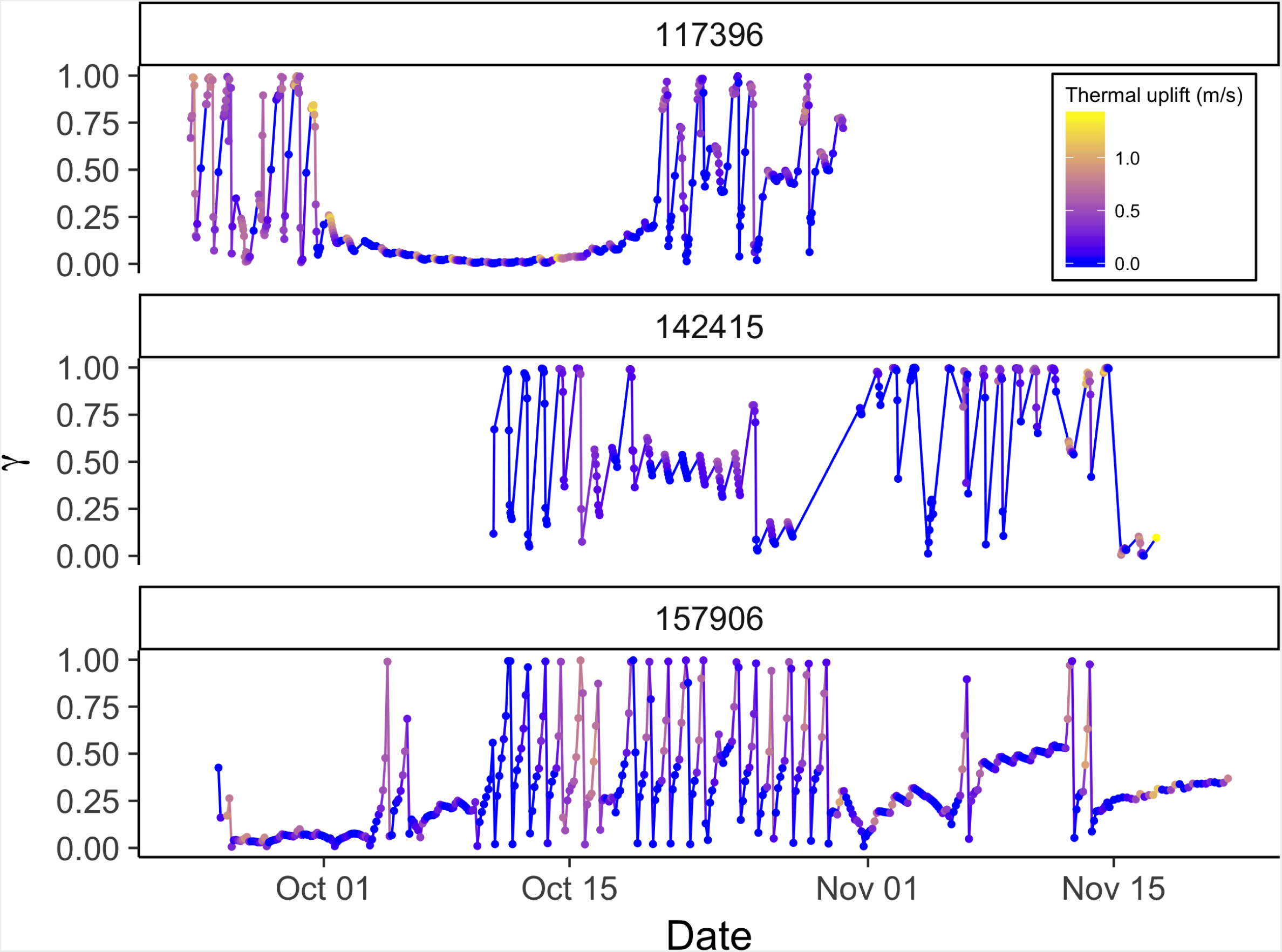
Time series of behavior parameter *γ* from correlated random walk model with full behavioral process (orographic uplift, thermal uplift, and tailwind support as predictors) for three golden eagles during fall migration with PTTs reporting on different duty cycles. Upper panel is 13 hourly centered on solar noon plus one at midnight, middle panel is 8 hourly centered on solar noon, and lower panel is fixed 3-hr interval. *γ* close to one reflect movements associated with migratory behavior, and *γ* close to zero stopover behavior. Points are times of observations, and lines are linear interpolations between points. Hue indicates intensity of thermal uplift, with yellow indicating greater thermal uplift and blue lower. Note the daily rhythm in behavior and extended stopovers as well as periods intermediate values suggesting fly-and-forage.

**Figure 3.**
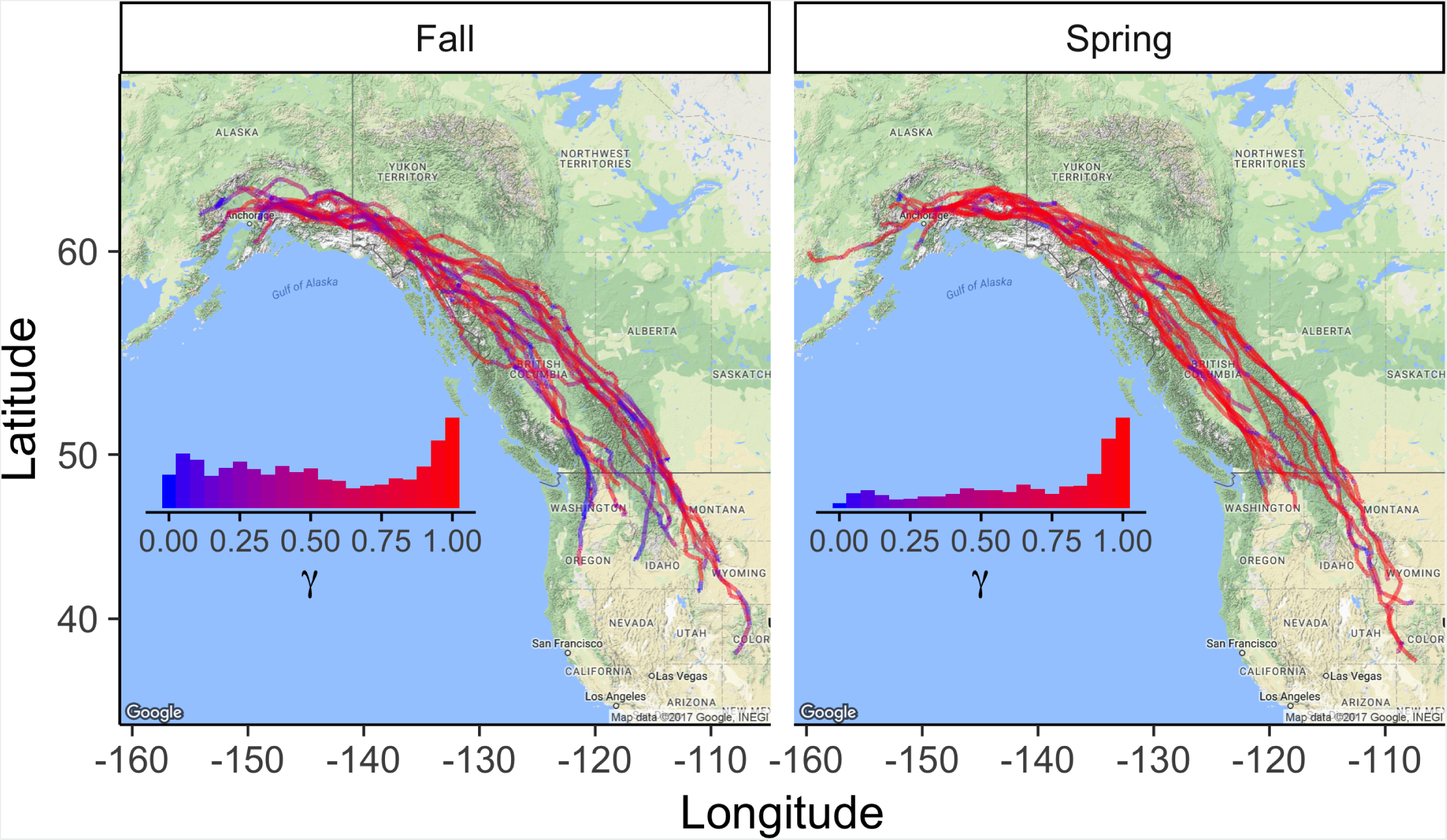
Golden eagle migration trajectories (*N* = 15 spring and *N* = 16 fall). Hue indicates value of behavioral parameter *γ* estimated with the correlated random walk model with full behavioral process, including orographic uplift, thermal uplift, and tailwind support as predictors. Insets show the relative frequencies of estimates of *γ* assigned to the displacements between observed daytime GPS locations. *γ* close to one reflect movements associated with migratory behavior, and *γ* close to zero stopover behavior. Daily rhythms, revealed in figures 1 and 2, are not apparent here because the birds moved so little at night.

There was also a clear effect of season on movement patterns and behavior. Spring was characterized by straighter, more direct trajectories and punctuated by slower, more tortuous, stopover-like movement; whereas, fall movements were much more tortuous overall and regular patterns in changes in movement rate and/or tortuosity less clear (Figs. 1–3). The distributions of estimated *γ* values also clearly indicate that daytime movements were most frequently directed migratory moves in the spring; whereas, in the fall, the bimodal distribution indicates more equivalent partitioning between directed migratory moves and slower stopover type movement, with significant time spent exhibiting behaviors associated with intermediate tortuosity and movement rate (Fig. 3).

### Environmental covariates

Including environmental covariates in the behavioral process improved model fit for almost all fitted migrations (Table 2). While there were differences in some environmental covariates between spring and fall (Table 1), parameter estimates from the full model (all covariates) indicate that there was little to no difference in effect of flight subsidies on behavior between spring and fall (Fig. 4, Table S1). Positive coefficients on the thermal uplift covariate indicate that increasing thermal uplift resulted in more highly-correlated displacements, or migratory movements. Despite that, there were some migration bouts not associated with great thermal uplift (Figs. 1 & 2). Coefficients close to zero for orographic uplift and tailwind indicate that, in general, they were not strong drivers of directionally-persistent movements, though they were probably occasionally used by birds for short movements when these subsidies were available.

**Table 2:**
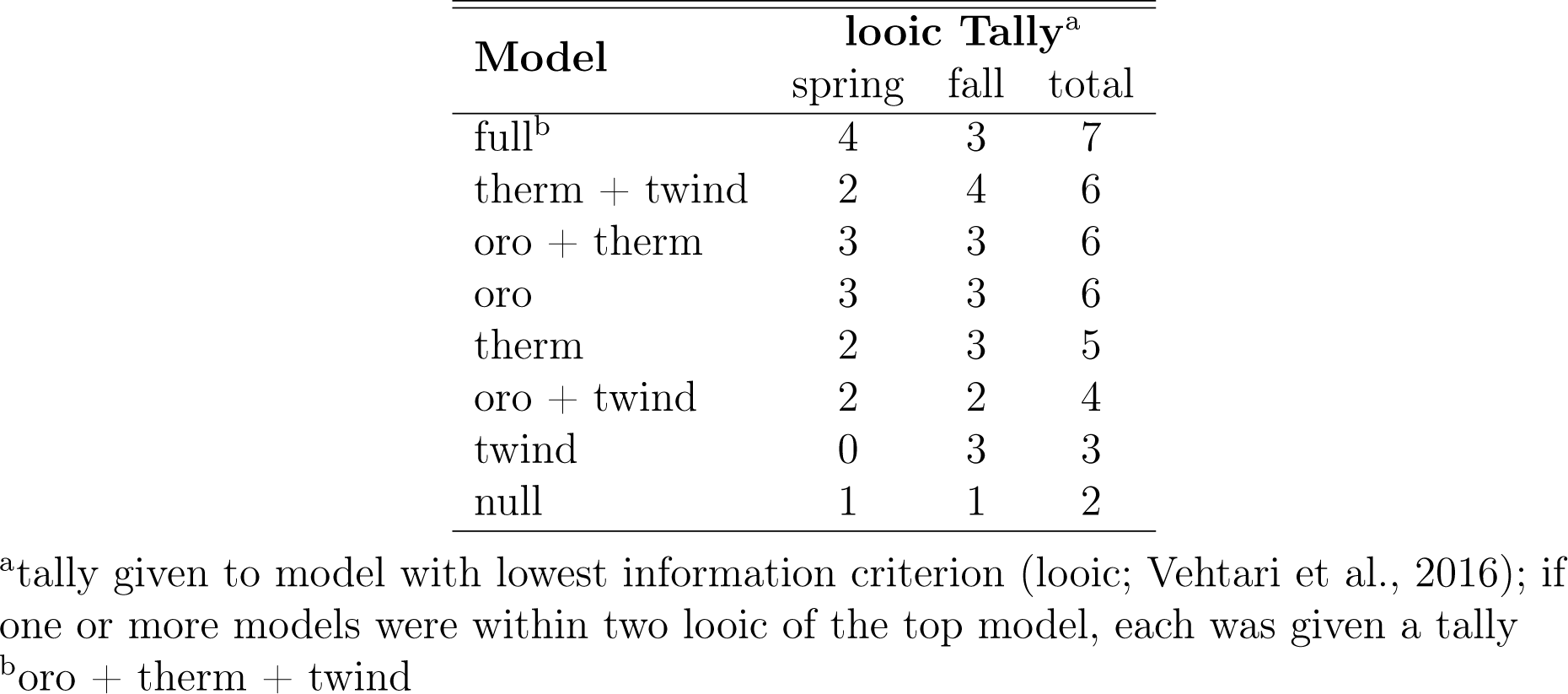
Number of golden eagle migration tracks recorded by GPS transmitters that each candidate formulation of the behavioral process in the correlated random walk model fit the best, according to approximate leave-one-out cross-validation (Table S1).

**Figure 4.**
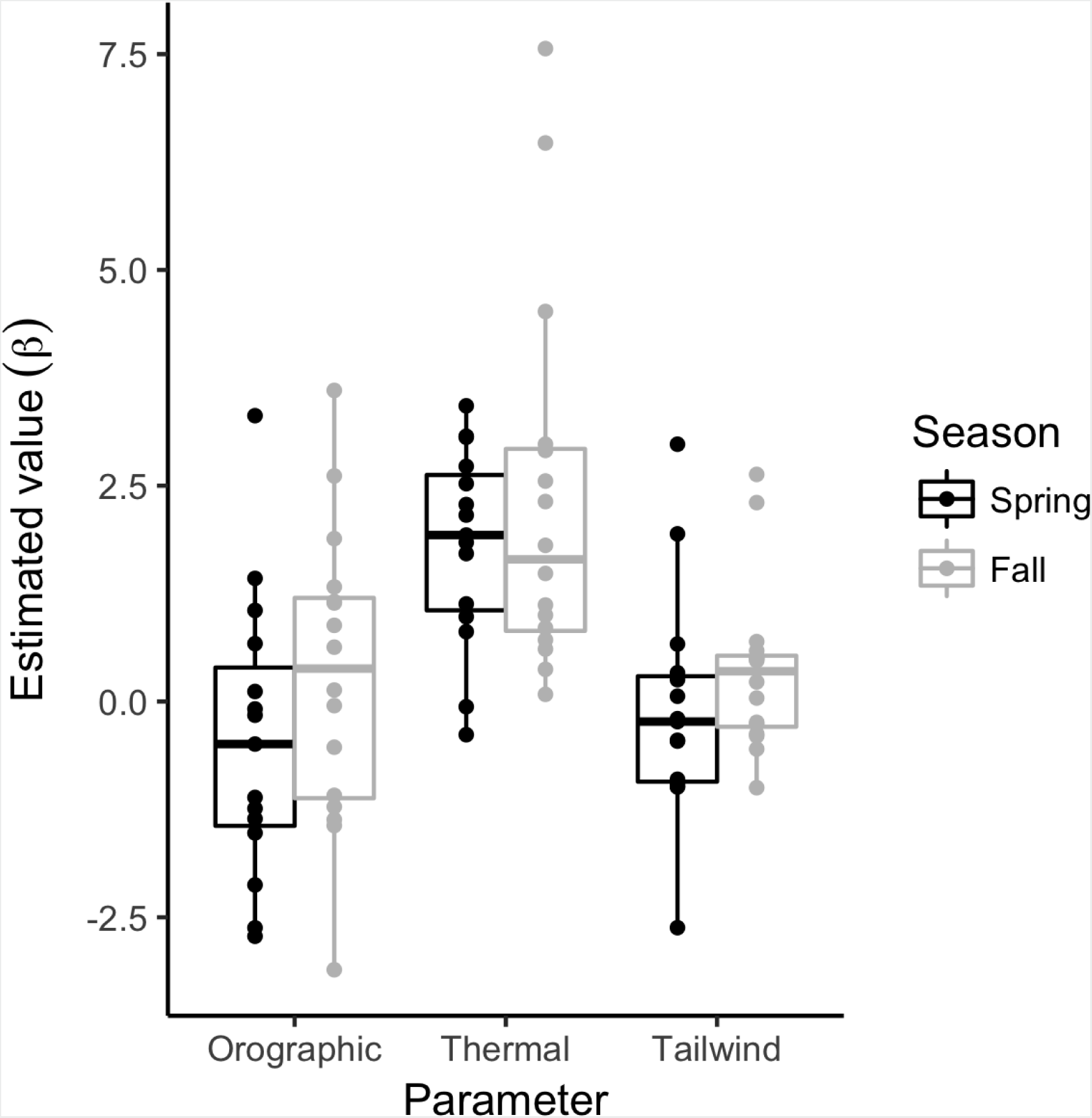
Point estimates of environmental covariate effect parameters (*β*_*ou*_, *β*_*tu*_, *β*_*tw*_) on golden eagle behavior and movements during migration (*N* = 15 spring and *N* = 16 fall). Estimates are from the correlated random walk model with full behavioral process, including orographic uplift, thermal uplift, and tailwind support as predictors.

Based on the model selection, the best-fitting formulation of the environmental drivers of the behavioral process was variable across individuals. However, in almost all cases, some form of flight subsidy was used and there was little difference between the spring and fall seasons in the pattern of subsidy use (Table 2). The high variability across individuals (Table S1) was likely due to differing weather patterns and thus subsidy sources encountered and/or used by each eagle as migrations were not synchronous (in time or space) across individuals. In addition, inter-individual variation was much larger than any difference attributable to demographic variables; we found no evidence that difference in sex nor age explained patterns of flight subsidy use during migration. Note, though, that all eagles included in this analysis were in adult plumage, so strong age effects would not necessarily be expected.

## Discussion

Here, we develop and demonstrate how dynamic parameter CRW models fit to GPS data reveal the effects of variable flight subsides available along migration routes. Use of these subsidies gives rise to diverse patterns in the movement of a long-distance soaring migrant. Behavioral changes occur continuously as available subsidies shift across space and time as migration proceeds. These key driving mechanisms underlie emergent movement paths, yet such processes are often hidden in the discrete satellite observations available. Our mechanistic modeling approach allowed linking of dynamic meteorology to changes in behavior, and those changes in behavior to the observed movement paths, revealing time series of behaviors more complex than individuals simply apportioning time between migration and stopover.

### Model performance

Incorporating time-varying parameters into movement models has been a relatively infrequently utilized approach (Breed et al., 2012; Auger-Méthé et al., 2017; Jonsen et al., 2019). Here we provide a case study for its utility and developed the approach for achieving practical biological inference about movement processes. Modeling the serial correlation in movement as a function of environmental covariates (equation 4), allowed simultaneous inference of behavior and the effect of environmental covariates on behavior from animal trajectories with regular and irregular duty cycles and containing missing observations. The behavioral patterns we found would be more difficult to reveal with a state-switching movement model (e.g. Hidden Markov Models (HMMs); Michelot et al., 2016) because each step would be forced into a discrete behavioral state from a set of usually 2-3 discrete states. Moreover, although hidden-state models have been introduced that have more than five discrete states (e.g., McClintock et al., 2012), these states can require ancillary data streams (e.g., accelerometry) to discriminate and remain extremely difficult to employ and interpret in practice (Patterson et al., 2017; Pohle et al., 2017). Finally, as HMMs include greater numbers of potential states, they tend to fit better than models with fewer states, even when additional states are not biologically meaningful nor sensible (Pohle et al., 2017). Implementing models with dynamic parameters that can be interpreted along a behavioral continuum is likely a much more natural approach for many animal movement questions.

#### Effects of tag programming

While our CRW model revealed the same patterns across duty cycles and was generally robust to the different duty cycles (Figs. 1 & 2), the most detail in daily behavioral rhythms was revealed in tracks with a fixed 3-hr time interval (lower panel in figure 2), as it provided data throughout the 24-hr day at regular intervals. The other duty cycles were initially chosen to minimize the risk of battery depletion overnight. Although generally robust, duty cycles did affect model fitting. HMC permitted Bayesian inference rather efficiently for our model, considering elevated correlation in the posterior of parameters due to the model formulation. Model fits typically took no more than a few hours, though tracks with much more than several hundred locations sometimes took longer. Preliminary fitting of our model with Stan and Template Model Builder (TMB; following Auger-Méthé et al. 2017) suggested that Maximum Likelihood estimation (when fit with TMB) tends to fail frequently when tag programming results in uneven temporal coverage of each day (e.g., our 2015 duty cycle), while Bayesian inference still provided sensible parameter estimates in most cases. Although the model presented herein and the model presented by Auger-Méthé et al. (2017) can make up for irregular time intervals between observations, they do have limitations. Breed et al. (2011) offer an in-depth discussion of tag programming and its effects on model fitting and inference.

### Flight subsidies as drivers migration of behavior

Thermal uplift is a flight subsidy dependent on daily atmospheric boundary layer dynamics, and it was clearly an important driver of the daily rhythm in eagle movement (Fig. 4, Table S1). Intense thermal uplift was often associated with the peaks in daily migration bouts (Fig. 1). The larger magnitude of the thermal uplift effect, relative to orographic uplift, was somewhat surprising, as many individuals in our sample followed the Rocky Mountains, a large potential source of orographic uplift. Golden eagles are known to use orographic uplift as a flight subsidy while migrating through the Appalachian Mountains in eastern North America (Katzner et al., 2015). Much of the Appalachians, however, is characterized by long, unbroken, linearly-oriented ridges. Wind blowing over these ridges produces long stretches of predictable orographic uplift (Rus et al., 2017). The Rocky Mountains, by contrast, are far more rugged and nonuniform, and conditions that might produce suitable upslope winds and mountain waves, as well as strong tailwinds, likely also generate violent turbulence and could impede efficient migratory flight. Soaring raptors have been shown to use small-scale turbulence to achieve subsidized flight (Allen et al., 1996; Mallon et al., 2016); however, unpredictable, nonstationary violent turbulence, which can occur in large, high-elevation mountain ranges (Ralph et al., 1997), could produce unfavorable migratory conditions. The large effect of thermal uplift, thus, could indicate that the Rocky Mountains, a spine that spans almost the entire migration corridor for this population, as well as some areas further west (Bedrosian et al., 2018), serves as a network of thermal streets for migrating eagles (Pennycuick, 1998). More explicitly, intense sun on south facing slopes would be expected to generate linear series of thermals that birds could glide between during both spring and fall migration. It is important to keep in mind that the migrants could capitalize on fine-scale, localized features of certain flight subsidies, like orographic uplift and tailwind, that may not have been captured by the interpolated meteorological data used in our analyses. However, model selection for models including those variables did indicate they explained some variance in eagle movement, which we discuss further below.

Despite meteorological conditions along migration paths that differed between spring and fall and a stark difference between behavioral budgets, our results showed no clear difference in the use of flight subsidies between the spring and fall seasons (Fig. 4). This finding contrasts with season-specific effects of flight subsidies on golden eagle migration shown phenomenologically in eastern North America, where thermal uplift was shown to be the key subsidy in migratory performance during spring, while tailwind with some additional support from thermal uplift is most important in the fall (Duerr et al., 2015; Rus et al., 2017). Although our results indicate that eagles use similar flight subsidizing strategies in both seasons, consistent with the differences from the eastern population, the actual behaviors performed during spring and fall migrations differed considerably. In spring, eagles used subsidies to drive a migration that allows timely arrival on the breeding grounds, consistent with a time minimization strategy. In the fall, flight was subsidized to minimize net energy use, which emerged as a much more diverse behavioral repertoire during a slower fall migration (Fig. 3; Miller et al., 2016). The more rapid and direct flight punctuated by bouts of tortuous, stopover-like movement in the spring (Fig. 3), suggest eagles pause, refuel, and/or perhaps wait for better migration conditions. This aligns some with a net energy maximization strategy (Hedenström, 1993; Miller et al., 2016), despite the need for timely arrival on the breeding grounds to avoid fitness costs (Both and Visser, 2001).

The behavioral time series of spring migrations showed some evidence of individuals responding less to thermal uplift as latitude increased (Fig. 1). This likely corresponded to a general decay in thermal uplift as latitude increased across individuals (Supplementary Material, Fig. S2). Reduced thermal uplift availability would be expected at higher latitudes due to the larger amounts of remaining spring snowpack and lower solar angles. Thus, golden eagles, and likely other soaring birds, migrating to high latitudes may need to budget behaviors carefully between time minimization and net energy maximization during spring migration to best take advantage of the reduced flight subsidy from thermal uplift and mitigate the greater energy demands of flight at higher latitudes.

While our results show that thermal uplift is the most important flight subsidy for the majority of migrating eagles sampled, the model selection indicated orographic uplift and tailwind had power in explaining eagle movement. Additionally, variability in top models across individuals (Tables 2 & S1) suggests among-individual variance in flight-subsidizing strategy. Although some of this variability can be attributed to real individual differences in behavioral strategy, it is at least as likely that individuals encountered different subsidies en route and used the subsidies they had available, as migrations across our sample were not synchronous. Given that orographic uplift and tailwind support parameter estimates were negative or close to zero for many individuals (Fig. 4, Table S1), those covariates likely predicted the periods of slower, more tortuous movements (i.e. *γ*_*i*_ closer to zero). Tailwind support occurred in top models for more individuals in the fall (Tables 2 & S1), which is consistent with findings from others (McIntyre et al., 2008; Rus et al., 2017) and suggests it may be important during southbound migrations. Finally, although there was variance among the types and combinations of subsidies used, the null model (without flight subsidies) was the best fitting model for very few tracks (Tables 2 & S1), evidence that weather and flight subsidies are of great importance to migrating golden eagles, and likely also to the migrations of similar soaring species.

### Daily rhythm & complex stopover

The full movement model revealed two clear, nested behavioral patterns in the long-distance migrations of golden eagles. First, there was a daily rhythm where inferred directed migratory movements (i.e. *γ*_*i*_ close to one) occurred most frequently around midday or early afternoon (Figs. 1 & 2). Mechanistic models of animal movement have revealed diel behavioral rhythms in other taxa (Jonsen et al., 2006). The basic aspects of daily rhythms in vertebrate behavior have hormone controls (Cassone, 1990), but the benefits can include balancing migration progress and foraging bouts (Newton, 2008). Soaring migrants also benefit by synchronizing diel movement patterns with diel atmospheric cycles. That is, consistent with our results, diurnal soaring migrants express a general circadian behavioral rhythm, where flight performance and behavior is strongly tied to thermal development of the planetary boundary layer to take best advantage of atmospherically generated flight subsides (Kerlinger et al., 1985; Leshem and Yom-Tov, 1989; Spaar and Bruderer, 1996, 1997; Mateos-Rodríguez and Liechti, 2012).

The second behavioral pattern revealed was a general stopover pattern, whereby eagles changed behavior for one to several days while en route (Figs. 1–3). These changes were consistent with searching movements (i.e. *γ* intermediate or close to zero), possibly representing foraging behavior. In terms of soaring raptors, however, very few reports of movement patterns and behavior during stopovers have been published. Stopover segments have been previously identified by speed or some other metric calculated from tracks, then excluded from subsequent analyses (e.g., Vansteelant et al., 2015; Katzner et al., 2015); occasionally, authors noted apparent enhanced tortuosity but explored it no further (e.g., Vansteelant et al., 2017). On occasions where stopover behavior was considered, classifications based on stay duration and travel distance or speed with hard, often arbitrarily chosen cutoffs between migrating and stopover segments were used (Duerr et al., 2015; Chevallier et al., 2011; Katzner et al., 2012; Miller et al., 2016). In contrast, our modeling framework aligned with the movement ecology paradigm (Nathan et al., 2008); it used the observed data—GPS locations, rather than a derived metric—and a theoretical movement process to infer behavior from movement patterns along tracks on a spectrum ranging from stopped to rapid, directionally-persistent movement.

Our analyses, however, showed that eagles still tended to continue along their migration route during periods of movement most resembling stopover, but with reduced movement rate and directional persistence (Figs. 1–3). This pattern suggests a joint migration/opportunistic foraging behavior that resembles fly-and-forage migration (Strandberg and Alerstam, 2007; Åkesson et al., 2012; Klaassen et al., 2017), which is consistent with observations of en route hunting behavior of golden eagles by Dekker (1985). Such behavior could be used to maintain balance between time expenses and energy intake, as it allows simultaneous migration progress and foraging.

This pattern does not fall very well within the “stopover” paradigm (Gill, 2007; Newton, 2008), however, as true stops during the migrations we observed were rare, except for expected nightly stops. Rather, migrants seemed to change their *pace*—either by slowing down, moving more tortuously, or both—but still generally moved toward their migratory destination (Figs. 1–3). Thus, instead of a discrete behavioral framework, whereby migrants switch between two migratory phases (migration and stopover) with very different movement and behavioral properties, we propose that for certain taxa, including some and perhaps many soaring migrants, as well some migrating species in other fluid environments such as fishes and marine mammals, a continuous alternative framework “migratory pacing” may be more appropriate and a natural way to interpret en route migratory behavior and movement dynamics. Soaring birds, even when energy reserves are relatively depleted, likely can still make steady progress toward a migratory goal when flight subsidies are available. Flapping migrants, on the other hand, would not be able to achieve this as readily due in part to the greater energy demands for sustained flight, and would require more regular refueling stopovers where migration progress is temporarily completely arrested. Both opportunism in foraging and use of energetic subsidies are likely key characters of fly-and-forage behavior and the ability to change pace of migration without actually stopping, as they relax the need for individuals to stopover in specific, food-rich habitats, which are required by most migrants with less flexibility in food and that lack the morphological specialization to maximally exploit the energetic subsidies available in moving fluids (Piersma, 2007; Gill, 2007).

Our model results revealed seasonal variability in migratory pacing by golden eagles. The tendency for eagles to exhibit movements matching fly-and-forage behavior, and pace their migrations more slowly was most apparent during fall migration. In contrast, spring migration was usually composed of much more punctuated events of slower-paced movements but these were still extended over space (Fig. 3), indicating the eagles pace their migration and employ a mixed behavioral strategy to some extent in spring as well. During spring, hibernating mammalian prey would be minimally available, leaving carrion, along with a few non-migratory and -hibernating species (e.g., ptarmigan *Lagopus* spp. and hare *Lepus* spp.), as major food sources, which could help explain the more punctuated bouts of slower-pacing. Alternatively, individuals could have been slowed by poor weather conditions (Rus et al., 2017). Scavenging large ungulate carcasses would be extremely rewarding in terms of energy accumulation. Much of the carrion we used successfully to capture eagles was large ungulate (e.g., moose *Alces alces*), strongly suggesting that the population we sampled uses carcasses during migration. The bimodal distribution for the behavioral parameter *γ* in fall shows that eagles tended to budget daytime behaviors approximately equally between rapid, directed and slower-paced movements (Fig. 3); the high frequency and range of intermediate values are, again, evidence for the more complex fly-and-forage and pacing dynamic, rather than eagles simply switching between stopover and migration. This behavioral complexity might be biologically important, allowing eagles to arrive on winter home ranges in better condition compared to migration strategies that do not incorporate en route foraging opportunity. In contrast to fall, daytime movements in the spring were typically faster-paced (i.e. larger-scale and directionally-persistent; Fig. 3), consistent with a time minimization strategy, where eagles need to partition time more in favor of migration progress to ensure timely arrival on breeding grounds (Hedenström, 1993; Alerstam, 2011; Miller et al., 2016). We thus see in eagles, and propose more generally, that such pacing varies between and within seasons along the continuum between time minimization and net energy maximization strategies (Alerstam, 2011; Miller et al., 2016). A migrant’s pace would be expected to depend upon their energetic demands, energetic subsidies available from the environment, and the importance of arriving at the migration terminus in a timely fashion (Nathan et al., 2008).

### Implications & conclusions

We developed and applied a movement model with time-varying parameters to help reveal the mechanisms underlying the migration of a long-distance soaring migrant that relies on incredibly dynamic flight subsidies. We found that variation in flight subsidies gives rise to changes in migrant behavior with thermal uplift seemingly most important. While these findings might be expected given previous phenomenological work (e.g., Duerr et al., 2012; Lanzone et al., 2012; Katzner et al., 2015; Vansteelant et al., 2015; Miller et al., 2016; Shamoun-Baranes et al., 2016; Rus et al., 2017), we were able to show how meteorology is a mechanism driving changes in movement patterns and thus behavior.

In the behavioral budgets of migrating golden eagles, we identified an expected daily rhythm, as well as evidence for behavioral dynamics that would allow nearly simultaneous foraging and migration, which is greater complexity than the traditional stopover paradigm allows. Migratory pacing, facilitated by fly-and-forage behavior, expands the traditional notion of stopover, whereby a bird migrates until resting and refueling is required, at which point it stops for a brief period in specific habitat suitable for efficient foraging (Gill, 2007; Newton, 2008). This advance was enabled by incorporating time-varying parameters into the movement model, which revealed new behavioral patterns during migration of long-distance soaring migrants. While time-varying, dynamic parameters have been infrequently employed in movement modeling (Breed et al., 2012; Jonsen et al., 2013; Auger-Méthé et al., 2017), we have shown it is a powerful approach that can overcome certain limitations in discrete state-switching models and help provide novel insight into animal behavior.

## Data accessibility

All movement data used for this manuscript are managed in the online repository Move-bank (https://www.movebank.org/; IDs 17680093 and 19389828). The data contain information considered sensitive by the State of Alaska, but they could be made available at the discretion of the Alaska Department of Fish & Game and U.S. Fish & Wildlife Service. Code to fit the movement model to data and example data are provided as supplementary material.

## Ethical approval

Field procedures were conducted following the ADF&G Animal Care & Use Committee protocol #2013-036 and University of Alaska Fairbanks Institutional Animal Care & Use Committee protocol #859448.

## Author contributions

JME and GAB conceived the ideas of the research presented herein. JME, MA-M, and GAB designed analytical methods, and TLB, CPB, and SBL designed the field methodology. TLB, CPB, SBL, and JME collected the data. JME analyzed the data, and led the manuscript. All authors contributed to drafts and editing of the manuscript and provided final approval for publication.

## Supporting information

example data

## Acknowledgements

We thank M. Kohan, B. Robinson, T. & D. Hawkins and many others for support in the field and J. Liguori and N. Paprocki for help aging eagles. We also thank T. Avgar, P. Doak, T. Katzner, K. Kielland, and C. McIntyre for helpful comments on drafts of the manuscript. Funding was provided by the Alaska Department of Fish & Game (ADF&G) through the federal State Wildlife Grant Program, and the U.S. Fish & Wildlife Service (USFWS) provided PTTs and data. JME was supported by the Calvin J. Lensink Fund during part of the project. MA-M acknowledges the support of the Natural Sciences and Engineering Research Council of Canada. The findings and conclusions of this paper are those of the authors and do not necessarily represent the views of the USFWS. This manuscript has been released as a pre-print at (Eisaguirre et al., 2019, https://www.biorxiv.org/content/10.1101/465427v2).

## Appendix 1: supplementary tables & figures

**Table S1:** Estimates of environmental covariate effects on golden eagle behavior and movements during migrations. Parameter estimates are from the correlated random walk model with full behavioral process, including and intercept (*β*_0_) orographic uplift (*β*_*ou*_), thermal uplift (*β*_*tu*_), and tailwind support (*β*_*tw*_) as predictors. Top model is the best fitting candidate of the behavioral process resulting from an approximate leave-one-out cross-validation model selection procedure.

| ID<sup>a</sup> | Season | Parameter<sup>b</sup> |  |  |  | Top Model<sup>c</sup> |
| --- | --- | --- | --- | --- | --- | --- |
| | | $\beta_0$ | $\beta_{ou}$ | $\beta_{tu}$ | $\beta_{tw}$ | |
| 117394 | spring | -2.68 | -1.52 | 0.98 | 0.67 | oro |
| 117396 | spring | -2.70 | -0.49 | 3.07 | -0.46 | - |
|  | fall | -0.53 | -3.11 | 0.08 | 0.49 | full<sup>d</sup> / oro + twind |
| 117397 | spring | -4.80 | 1.43 | 3.06 | 0.25 | therm + twind / therm |
| 117398 | spring | -2.21 | -0.09 | 0.81 | -0.45 | oro / therm |
| 135411 | spring | -2.55 | -2.62 | 1.84 | -0.98 | oro + therm / full / oro + twind |
| 135413 | spring | -2.98 | 0.12 | 1.13 | 0.06 | oro |
| 135414 | spring | -0.74 | -1.11 | -0.38 | 1.94 | oro + twind |
| 135417 | spring | -2.53 | -1.36 | 2.28 | -0.20 | oro + therm |
|  | fall | -2.90 | -0.53 | 0.85 | 2.30 | therm |
| 135421 | fall | -2.78 | -1.09 | 2.91 | 0.69 | twind |
| 135423 | spring | -2.80 | -1.24 | 2.52 | -0.99 | null |
|  | fall | -2.12 | 0.13 | 1.00 | -0.24 | therm + twind |
| 135425 | fall | -2.29 | -1.37 | 1.81 | 0.51 | twind |
| 142404 | spring | -2.90 | 3.31 | 1.71 | -2.62 | full |
| 142405 | spring | 2.34 | 1.06 | 1.93 | -0.96 | full / oro + therm |
| 142407 | spring | -2.11 | -0.16 | 2.72 | 2.98 | therm + twind / full |
|  | fall | -0.57 | -1.22 | 0.37 | 0.23 | full / oro + therm |
| 142408 | spring | -2.66 | -2.13 | -0.06 | -0.90 | - |
|  | fall | -4.82 | 2.61 | 2.32 | -1.00 | oro + therm |
| 142409 | spring | -1.29 | 0.67 | 2.16 | 0.33 | - |
| 142415 | fall | -4.26 | 1.16 | 0.61 | 2.63 | oro + twind |
| 142417 | spring | -0.83 | -2.72 | 3.43 | -0.23 | - |
| 157895 | fall | -0.65 | 0.88 | 1.12 | 0.04 | therm / oro |
| 157896 | fall | -0.73 | 0.63 | 1.48 | -0.40 | - |
| 157899 | fall | -2.10 | 1.89 | 6.47 | 0.47 | oro / therm + twind / therm |
| 157900 | fall | -0.42 | -0.05 | 2.98 | -0.27 | null / therm + twind |
| 157902 | fall | -1.98 | -1.44 | 7.56 | -0.38 | oro + therm |
| 157903 | fall | -0.74 | 1.14 | 0.71 | 0.59 | full |
| 157904 | fall | -1.31 | 3.60 | 2.55 | 0.51 | oro / oro + therm |
| 157906 | fall | -1.16 | 1.33 | 4.52 | -0.55 | twind |
| *mean* | spring | -2.10 | -0.46 | 1.81 | -0.10 |  |
|  | fall | -1.83 | 0.29 | 2.33 | 0.35 |  |
| *sd* | spring | 1.57 | 1.64 | 1.13 | 1.31 |  |
|  | fall | 1.34 | 1.72 | 2.18 | 0.96 |  |

<sup>a</sup>individual eagles; blank indicates second season for previous ID
<sup>b</sup>posterior means of parameter estimates
<sup>c</sup>model with lowest looic (Vehtari et al., 2016); all models within two looic of the top model listed in order of fit (separated by /); - indicates null did not converge
<sup>d</sup>oro + therm + twind

**Figure S1:**
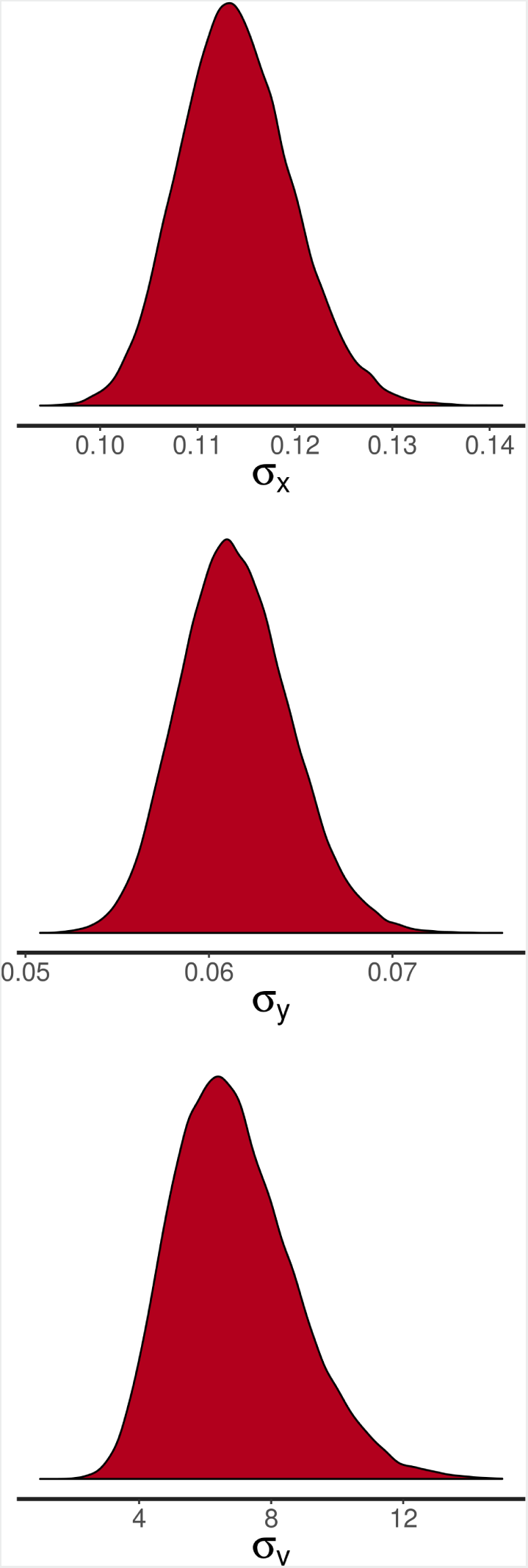
Posterior plots of variance components of the correlated random walk model with orographic uplift, thermal uplift, and tailwind support as behavioral predictors for the spring track of golden eagle 135423. Curves are approximately Gaussian, indicating the model was well behaved and likely converged to the posterior.

**Figure S2:**
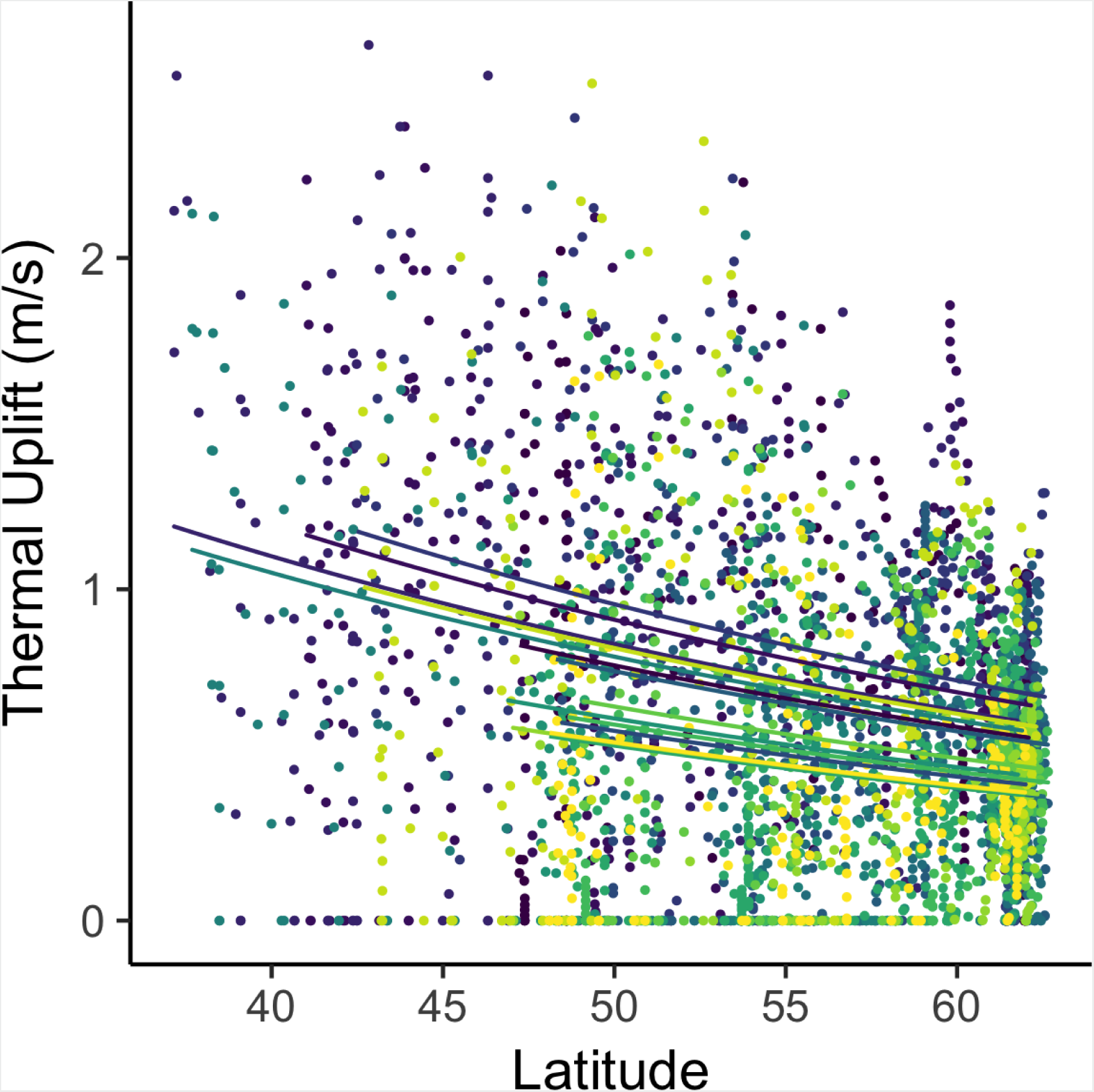
Interpolated thermal uplift as a function of latitude during spring migration. Hue corresponds to individual. Curves are from the individual level of a Bayesian hierarchical Gamma regression. The 95% Bayesian credible interval for the latitude coefficient was –0.040 *< β*_*lat*_ *<* –0.015, strong evidence for a decreasing trend in thermal uplift with increasing latitude.

## Appendix 2: code

### Stan model

~~~
data {

  int N;             // # of fixes in track
  vector[N] x;       // x coordinates
  vector[N] y;       // y coordinates
  vector[N] dt;      // time intervals

  vector[N] oro;     // covariates
  vector[N] therm;
  vector[N] twind;
  vector[N] tod;

}

transformed data {

  vector[N] c_oro;
  vector[N] c_therm;
  vector[N] c_twind;
  vector[N] oro_inter;
  vector[N] therm_inter;
  vector[N] twind_inter;

  // shifted log transform and standardize
  c_oro = (log(oro + 1) * (0.5 / sd(log(oro + 1))));

  c_therm = (log(therm + 1) * (0.5 / sd(log(therm + 1))));
  // center and standardize twind
  c_twind = ((twind) * (0.5 / sd(twind)))
  - mean(twind * (0.5 / sd(twind)));

  // interactions
  oro_inter = c_oro .* tod;
  therm_inter = c_therm .* tod;
  twind_inter = c_twind .* tod;

}

parameters {

  vector[N] gamma_raw;                   // logit behavior parameter
                                         //-- time-varying, correlates steps
  real<lower=0> sigmax;    // variance in x
  real<lower=0> sigmay;    // variance in y
  real<lower=0> sigmav;    // behavior variance
  vector[4] beta;                        // covariate coefficients

}

transformed parameters{

  // introduce logit link
  vector<lower=0,upper=1>[N] gamma;

  for(j in 1:N){
  gamma[j] = inv_logit(gamma_raw[j]);
 }

}

model {

  // NOTE: Stan uses st. dev. for normals, whereas JAGS uses precision
  // Stan also truncates appropriately based on specified constraints
 

  // prior on behavior process noise
  // prior density away from zero--assume there is variability in behavior
   sigmav ∼ normal(3 , 3);

  // priors on movement process noise--close to zero
  sigmax ∼ normal(0 , 1);
  sigmay ∼ normal(0 , 1);

  // priors on coefficients
  // prior density on zero--assume no effect of covariates on behavior
  beta ∼ student_t(5, 0 , 2.5);

  for (i in 3:N) {

     // behavior linear combination of covariates plus previous behavior
     // this can be modified to make candidate formulations
     gamma_raw[i] ∼ normal(gamma_raw[i-1] + beta[1]
     + beta[2] * oro_inter[i] + beta[3] * therm_inter[i]
     + beta[4] * twind_inter[i] , dt[i] * sigmav);

    // movement process is independent in x and y
    x[i] ∼ normal(x[i-1] + gamma[i] * (dt[i] / dt[i-1])
    * (x[i-1] - x[i-2]) , dt[i] * sigmax);

    y[i] ∼ normal(y[i-1] + gamma[i] * (dt[i] / dt[i-1])
    * (y[i-1] - y[i-2]) , dt[i] * sigmay);

}

}

generated quantities {

  // generate log likelihoods for PSIS-LOO
  // test prediction of next step from previous with estimated gamma
  vector [N] log_lik;
  log_lik[1] = 0.1; // need something here; simplest to fix across tracks
  log_lik[2] = 0.1;

  for (n in 3:N){

     log_lik[n] = normal_lpdf(x[n] | x[n-1] + gamma[n]
     * (dt[n] / dt[n-1]) * (x[n-1] - x[n-2]) , dt[n] * sigmax)
     *normal_lpdf(y[n] | y[n-1] + gamma[n] * (dt[n] / dt[n-1])
     * (y[n-1] - y[n-2]) , dt[n] * sigmay);

}

}
~~~

### R code

~~~
library(rstan)
library(loo)

#################################
## model variables
# x -- vector of x coordinates
# y -- vector of y coordinates
# dt -- vector of time intervals
# N -- number of fixes in track
# oro -- vector of raw orographic uplift data
# therm -- vector of raw thermal uplift data
# twind -- vector of raw tailwind data
# tod -- vector of times of day (1=day, 0=night)
################################

dat = read.csv(‘examp_dat.csv’)

x = dat$x
y = dat$y
dt = dat$dt
N = nrow(dat)
oro = dat$oro
therm = dat$therm
twind = dat$twind
tod = dat$tod

# fit Stan model with HMC, using default no-u-turn sampler
# model code should be saved as a .stan file
# arguments should be modified as required
stan.fit = stan(“model.stan”,
         data = list(x, y, dt, N, oro, therm, twind, tod),
         chains = 2,
         iter = 1000,
         thin=1,
         cores = 1,
         control = list(adapt_delta = 0.9))
# last argument helps with divergent transitions

# PSIS-LOO
log_lik = extract_log_lik(stan.fit)
loo = loo(log_lik_00)
print(loo)
# use compare(loo1, loo2, …) to rank candidate models
~~~

